# Skin biomarkers for cystic fibrosis: a potential noninvasive approach for patient screening

**DOI:** 10.1101/182733

**Authors:** Cibele Zanardi Esteves, Letícia de Aguiar Dias, Estela de Oliveira Lima, Diogo Noin de Oliveira, Carlos Fernando Odir Rodrigues Melo, Carla Cristina Souza Gomez, José Dirceu Ribeiro, Antônio Fernando Ribeiro, Carlos Emílio Levy, Rodrigo Ramos Catharino

## Abstract

**Background:** Cystic fibrosis is a disabling genetic disease with an increased prevalence in populations with European heritage. Currently, the most used technique for collection of cystic fibrosis samples and diagnosis is provided through uncomfortable tests, with uncertain results, mostly based on chloride concentration in sweat. Since cystic fibrosis mutation induces many metabolic changes in patients, exploring these alterations might be an alternative to visualize potential biomarkers that could be used as interesting tools for further diagnostic upgrade, prioritizing simplicity, low cost and quickness.

**Methods:** This contribution describes an accurate strategy to provide potential biomarkers related to cystic fibrosis, which may be understood as a potential tool for new diagnostic approaches and/or for monitoring disease evolution. Therefore, the present proposal consists of using skin imprints on silica plates as a way of sample collection, followed by direct-infusion high-resolution mass spectrometry and multivariate data analysis, intending to identify metabolic changes in skin composition of cystic fibrosis patients.

**Results:** Metabolomics analysis allowed identifying chemical markers that can be traced back to cystic fibrosis in patients’ skin imprints, differently from control subjects. Seven chemical markers from several molecular classes were elected, represented by bile acids, a glutaric acid derivative, thyrotropin releasing hormone, an inflammatory mediator, a phosphatidic acid, and diacylglycerol isomers, all reflecting metabolic disturbances that occur due to of cystic fibrosis.

**Conclusion:** The comfortable method of sample collection combined with the identified set of biomarkers represent potential tools that open the range of possibilities to manage cystic fibrosis and follow the disease evolution. This exploratory approach points to new perspectives about cystic fibrosis management and maybe to further development of a new diagnostic assay based on them.

## 1. INTRODUCTION

Cystic Fibrosis (CF) is a recessive genetic exocrine disorder that affects the transport of fluids and electrolytes through cell membrane, resulting in abnormal viscous secretion in multiple organs. Although this disease affects all racial and ethnic groups, Caucasians are the most affected; 1 in every 2000-3000 European newborns presents CF ^1^. Soon after this disease was characterized, in 1953 ^2^, cystic fibrosis was considered almost inevitably lethal, mainly at the first decade of life. However, improvements at diagnostic methods and clinical management have led to a considerable increase in life expectancy for CF patients over the years, as shown by recent statistical data indicating that the birth cohort of the year 2000 may present a survival median of 50 years ^3^. Although most CF patients are able to manage well disease, their quality of life is still affected by limitations such as physiological functioning, endobronchial infection and pancreatic insufficiency ^4^.

All cystic fibrosis manifestations are caused by mutations affecting the cystic fibrosis transmembrane conductance regulator (CFTR). Although there are multiple mutations at CFTR gene associated with CF disorders, the most common corresponds to a deletion of a phenylalanine at position 508 on chromosome 7, called F508del mutation ^5^. More than 1000 CFTR mutations are known to cause CF; some may be responsible for milder CF phenotypes, while others present more severe manifestations. These differences at CFTR gene mutation locus generates great heterogeneity, making clinical and even laboratory diagnosis difficult tasks ^6^.

The two most used tests for CF laboratory diagnosis are DNA analysis and sweat test; the first one is applied to recognize known mutations on CFTR gene, however, gene mutations not reported yet may not be diagnosed due to a limited number of gene alterations present in mutation panels, predefined for commercial laboratory tests (e.g. 97 mutations) ^7^. Furthermore, this test allows the analysis of the complete CFTR gene sequence; nonetheless, the available current genetic analyses are unable to detect mutations that occur out of coding regions, which renders the assessment of these regions even more difficult. Moreover, it is an expensive and time consuming process that is performed in specialized molecular and genetics laboratories, making it a test with limited access for patients ^8^.

In addition to DNA analysis, the most widely employed laboratory diagnostic test in CF diagnosis is the quantitative pilocarpine iontophoresis testing (QPIT), also known as sweat test ^9^. Although it is an uncomfortable method, QPIT is considered the Gold standard test, and consists in the determination of chloride concentration in sweat; in this test, the normal upper limit can be <30 mmol/L for infants or <40 mmol/L for older adults ^10^. For sweat collection, perspiration is induced on skin of the forearm through local administration of pilocarpine nitrate, followed by the application of a battery-operated electric current that stimulates rapid and increased sweating rate. After sweat collection, chloride levels are evaluated and concentrations above 60 mmol/L are considered positive for Cystic fibrosis. ^9-11^.

Although QPIT is considered the method of choice for CF diagnosis, there is an obvious window of inaccurate diagnosis for chloride concentrations between 30 and 60 mmol/L, which classifies patients within this range as indeterminate cases, requiring further evaluation ^10^. Considering chloride quantification < 30 mmol/L as “normal” and ≥ 30 mmol/L as “at risk” for CF, QPIT sensitivity and specificity are 100% and 92.8%, respectively ^12^. However, the current standard diagnostic method does not differentiate positive and negative patients for CF who present chloride concentration between 30 mmol/L and 60 mmol/L. In addition to the large borderline window, the test procedures are liable of failure, mainly during steps such as sweat collection, weighting, dilution and elution, which hamper a more refined diagnosis. Furthermore, accurate sweat test results may be challenging taking into account the high frequency of technical errors and misinterpretation that may lead to types I and II errors ^13^. Therefore, development of new techniques is needed to improve CF diagnosis, reduce the borderline window and embrace the most cases of cystic fibrosis.

The metabolomics approach has been growing as a promising tool for monitoring biochemical pathways and their relationship with biological phenotypes in an integrated perspective ^14^. Metabolomics evaluation of biological samples allows researchers to identify biomarkers or sets of biomarkers that could be useful to differentiate health and disease profiles, which may lead to more accurate diagnostic tools ^15^. In this context, this contribution intended to provide a set of chemical markers for cystic fibrosis, based on a noninvasive sweat collection method, associated with high-resolution mass spectrometry. Finally, we aimed at providing some guidance with molecules that improve the knowledge about cystic fibrosis metabolism, and may work as alternative options for the development of new potential diagnostic tools.

## 2. METHODS

### 2.1 Patients’ selection

Sixteen patients were selected in a retrospective fashion at CF Clinic of the Clinics Hospital at University of Campinas. Inclusion criteria were that patients presented at least one sweat test with chloride concentration greater than or equal to 60 mEq/L, and presence of homozygosis for the mutation F508del, characterized by genetic studies. Exclusion criteria were the presence of skin lesions on the patients’ back, local of sample collection, and/or another skin disease. After selection, these patients were submitted to sample collection during scheduled visits to the hospital. The control (CT) group was composed by healthy individuals at the same age and gender as the CF group, recruited from schools and universities. Sample collection was approved by the Research Ethics Committee of the School of Medical Sciences – University of Campinas/Brazil (Protocol number: 1.100.978). All experiments were performed in accordance with relevant guidelines and regulations regarding samples of human origin.

### 2.2 Sample collection and preparation

Skin metabolites were collected from both CF and CT groups with plates of silica gel 60G, suitable for thin-layer chromatography (Merck, Darmstadt, Germany). These plates were cut into small squares of 1 cm^2^ and were superposed on the individuals’ back for 1 minute. Then, silica plates were extracted with 1000 μL of Methanol: Water (1:1), and 1 μL ammonium hydroxide was added to each sample to facilitate ionization.

### 2.3 Mass spectrometry

Samples were directly injected in an ESI-LTQ-XL Orbitrap Discovery instrument (Thermo Scientific, Bremen, Germany) with nominal resolution of 30,000 (FWHM) under the following conditions: flow rate of 10 ¼L.min^-1^, sheath gas at 10 arbitrary units, spray voltage of 5 kV and capillary temperature of 280°C. Analyses were performed in triplicates and all data were acquired in the negative ion mode at the mass range of 150-700 *m/z*.

### 2.4 Statistical analysis and biomarker election

An Orthogonal Partial Least Squares Discriminant Analysis (O-PLS-DA) using the online software MetaboAnalyst 3.0 (www.metaboanalyst.ca) ^16^ was performed to select characteristic ions from each group; the VIP score list (Variable Importance in Projection) generated by the software was considered for ions choice.

The online databases consulted were METLIN (Scripps Center for Metabolomics, La Jolla, CA), HMDB version 3.6 (Human Metabolome database – www.hmdb.ca), Lipid MAPS online database (University of California, San Diego, CA – http://www.lipidmaps.org), and KEGG Pathway Database (Kyoto Encyclopedia of Genes and Genomes – www.genome.jp/keeg/pathway.html). These databases were chosen to identify the potential biomarkers elected by statistical analysis, and it was established that mass error should not be greater than 2 ppm.

## 3. RESULTS

During the study, 16 CF patients were selected, with ages ranging from 5 to 19 years old (mean: 12 years; median: 13 years), with 56% males and 44% females. The sweat test presented an average concentration of chloride of 111 mEq/L (minimum: 76 mEq/L; maximum: 166 mEq/L). All patients presented deficits in lung function and 69% presented two or more different microorganism isolated from sputum within the past 12 months. Regarding further clinical conditions, 100% of patients present pancreatic insufficiency, 67% have evidence of liver disease, 44% make use of ursodeoxycholic acid and 19% developed diabetes. Regarding the control group, it was composed of 16 healthy individuals within the same gender and age range as the CF group and was submitted to sample collection at the same period as CF patients.

HRMS analysis on negative ion mode was performed, and Figure 1 shows representative mass spectra from each group. Information extracted from spectral data served as basis for the statistical analysis using O-PLS-DA, which compared the control and CF groups to elect biomarkers for CF. O-PLS-DA plot (Figure 2) shows complete separation of control and CF groups; from these analysis, 1 biomarker was selected for control group and 7 biomarkers for CF group, all presenting VIP scores above 2. Detailed information of the elected biomarkers for control and CF groups can be seen in Table 1.

**Figure 1.**
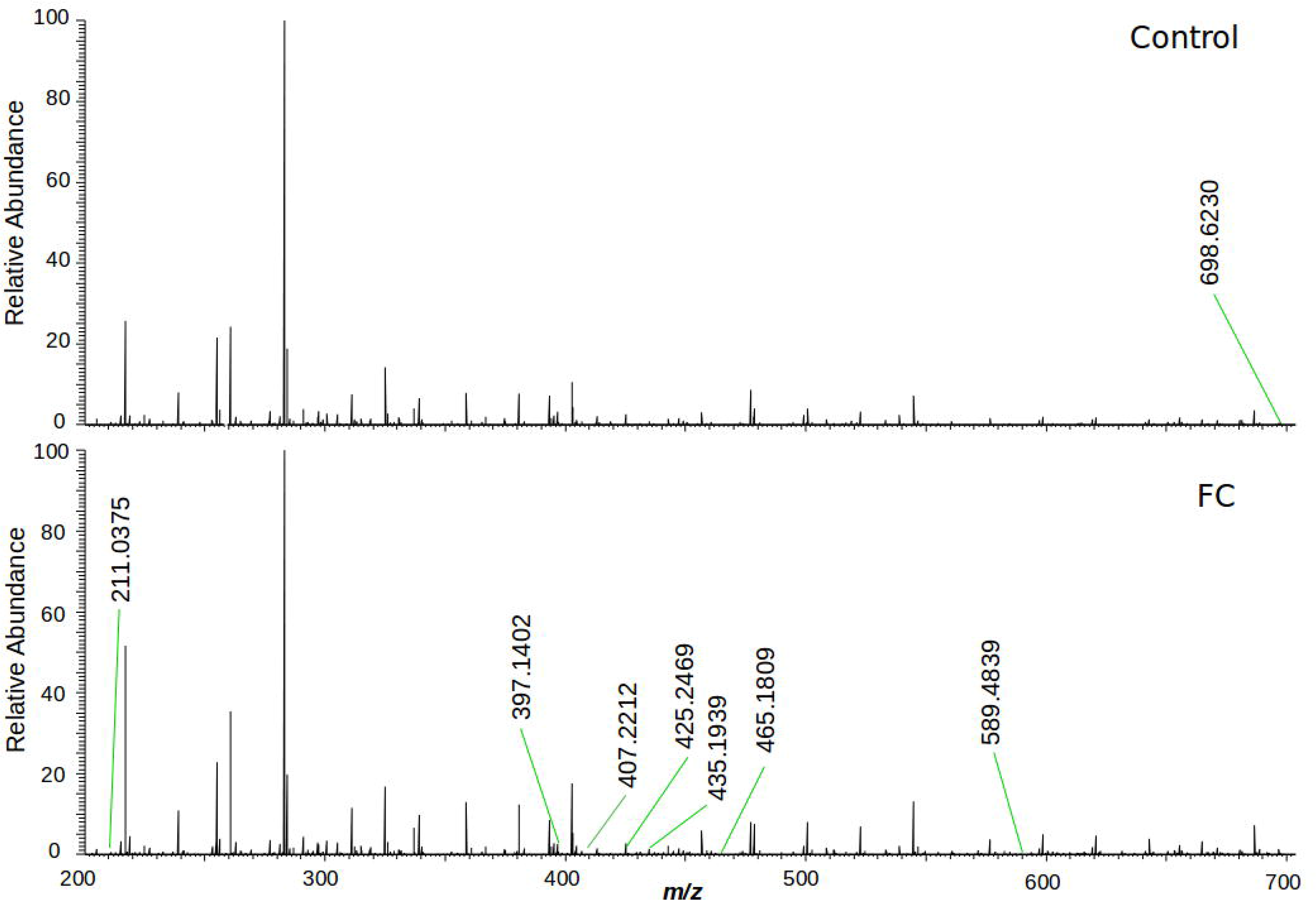
Representative mass spectra comparing the skin imprints of control individuals and FC patients. Negative ion mode.

**Figure 2.**
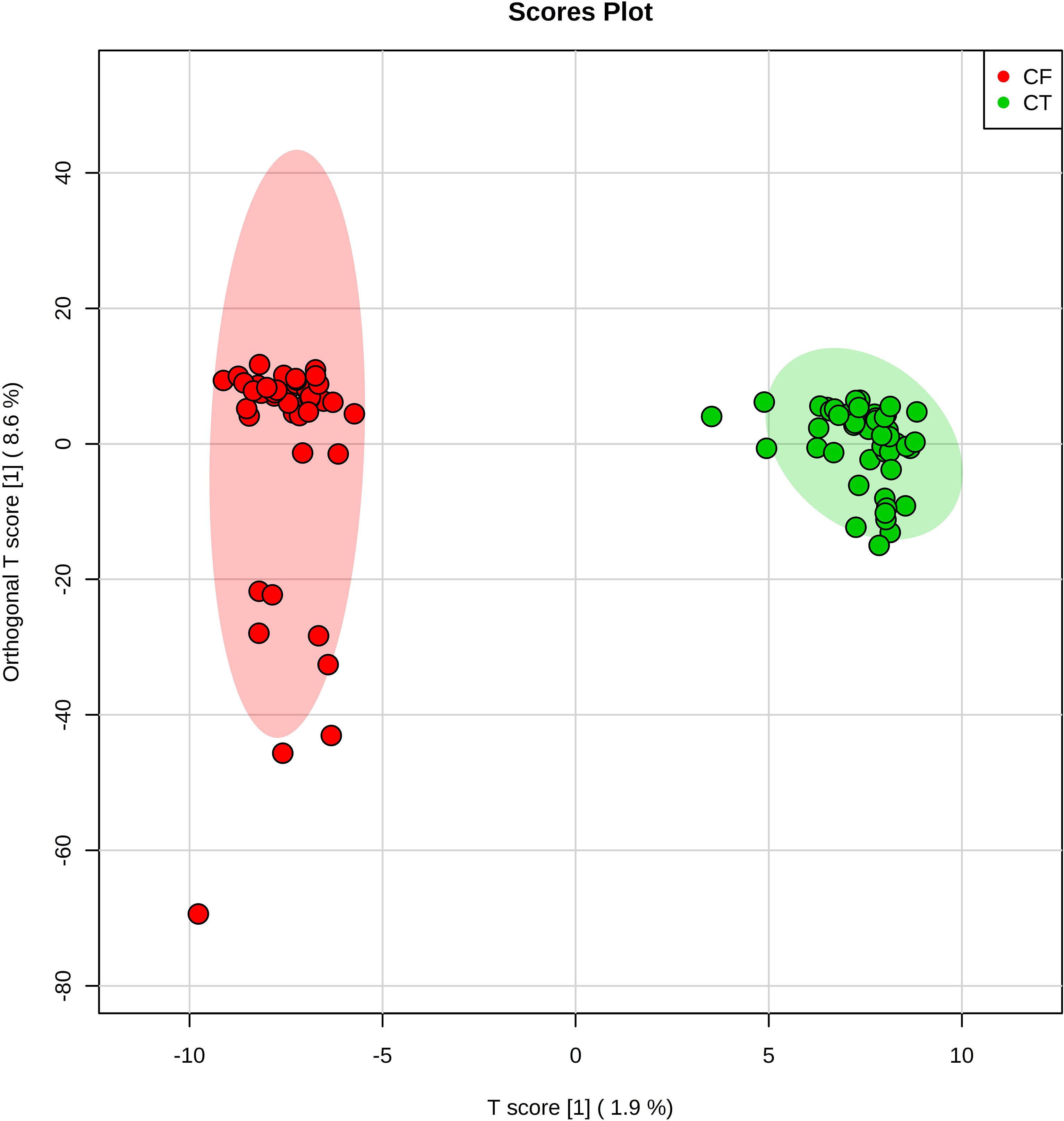
Two-dimensional plot provided by orthogonal partial least squares discriminant analysis (O-PLS-DA). It is possible to observe a clear separation between control subjects and patients with cystic fibrosis.

**Table 1.**
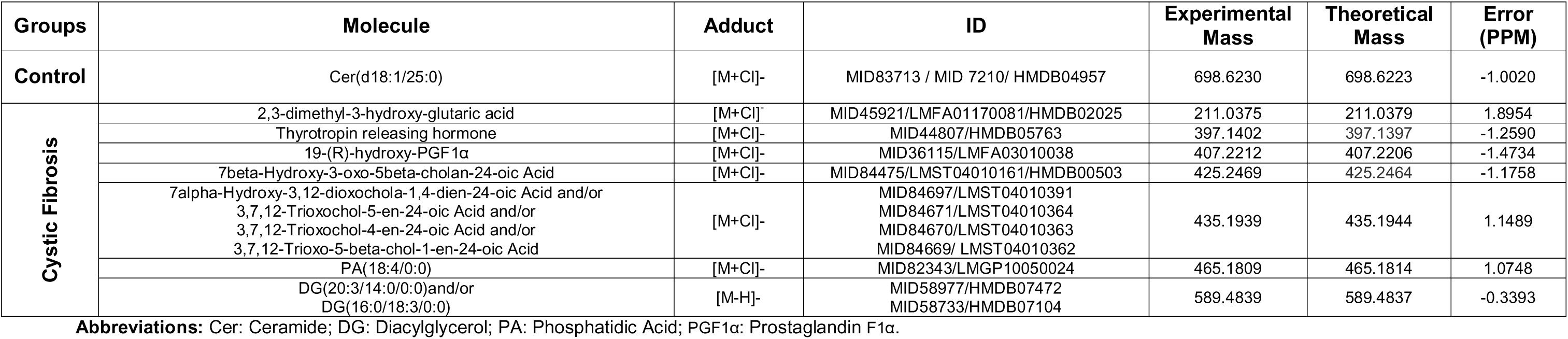
Biomarkers selected by O-PLS-DA, elucidated by HRMS. Negative ion mode.

## 4. DISCUSSION

Although sweat test remains the gold-standard method for CF diagnosis, there are some points that can be improved. The test lacks sensitivity and specificity at the concentration range of 30 – 60 mmol/L of chloride in sweat. In addition, sweat collection may be time-consuming and uncomfortable for the patient, especially for babies and children. Due to frequent methodological errors, retesting is often required more than once to present confirmatory results, which may be stressful considering the patients are, for the most part, children ^17^.

Intending to find chemical markers that could guide researchers through new methodologies, we have proposed a method based on mass spectrometry and metabolomics that presents a simple and quick workflow for sample collection, data acquiring and processing. For that, silica plates were chosen for sample collection due to its useful particularities of adsorption and stabilization of molecules, such as those present in the skin surface, what seems advantageous and dispenses the need for additional apparatus for sample preserving ^18^. This relatively simple sample collection and preparation, coupled with direct-infusion mass spectrometry, provides a more straightforward analysis protocol for biomarkers monitoring. With less sample preparation steps, the errors associated to the methodology decrease and the method becomes more reproducible, which is in accordance with a primary requisite to standardize a method in biomarkers identification: reproducibility.

### 4.1 Control group biomarker

A healthy skin is extremely important for humans, as it provides protection against environment injuries and covers the underlying tissues. It is composed by different layers: stratum corneum, epidermis, dermis and the fat layer. The outermost skin layer, stratum corneum, is composed by keratinocytes and lipids, specifically ceramides (50%), cholesterol (25%) and fatty acids (10 – 20%). These lipids provide fundamental limitation to water and electrolyte movement, and act as a barrier against microorganisms’ invasion ^19b^. Consequently, epidermis presents unique structures, such as epidermal ceramides that are not found in any other cell types in the human body ^19a^. For instance, the present study has identified Cer (d18:1/25:0) (698 *m/z*) in the epidermis of control subjects, an odd-numbered fatty acid carbon chain, uncommonly found in human cells in general, but present among epidermis’ lipids. About 30% of the total ceramide (Cer) present in the epidermis is represented by Cer containing odd-numbered fatty acids ^19a^, which corroborates the election of Cer (d18:1/25:0) as a control group marker. Therefore, this class of Cer stands out in this comparison, defining the control group’s skin, as determined by O-PLS-DA.

### 4.2 Cystic fibrosis biomarkers

Distinct molecules were identified through the present method as cystic fibrosis biomarkers, and each one is associated with metabolic dysfunctions observed in this disease. In healthy subjects, CFTR channel gating is predominantly regulated by protein kinase A (PKA) phosphorylation, which mediates gene expression, conformational changes, and protein trafficking from the endoplasmic reticulum (ER) to Golgi apparatus and cell surface ^20^. However, the phenylalanine deletion at position 508 (F508del) of the CFTR gene in cystic fibrosis reduces the rates of phosphorylation and may contribute to improper folding, defective trafficking, and slower rates of channel activation ^20-21^.

The F508del-CFTR is produced and retained in the ER, and undergoes ER-associated degradation through the proteasome pathway. Nevertheless, a limited number of misfolded proteins reach the Golgi apparatus for glycosylation and few of them reach cell surface ^20^. The export from ER depends on the interaction between the CFTR and the coat complex (COP) budding machinery ^20, 22^. Formation of COPII coated vesicles depends on the hydrolysis of phosphatidylcholine (PC) to phosphatidic acid (PA) by phospholipase D (PLD) ^22^. The relationship between PA and CFTR trafficking remains unknown, but Hashimoto and coworkers ^22^ showed that PLD-mediated PA formation is required for CFTR transport from ER to Golgi. In cystic fibrosis, it may be assumed that the organism is trying to maintain a compensatory mechanism of PA synthesis to enhance the concentration of COPII and the transport of F508del-CFTR to maturation in the Golgi apparatus, aiming to recover the imbalance caused by inoperative CFTR mutated. Therefore, the election of PA (465 *m/z*) as a CF biomarker in the present study is coherent with the cellular physiological condition. Considering PA’s importance for different intracellular events, different studies have also demonstrated PA as a regulatory lipid, even in signal transduction ^23a^. Accordingly, intracellular phosphatidic acid must be tightly regulated, which occurs through turnover into diacylglycerol (DG). Among the present results, diacylglycerol isomers (589 *m/z*) were also elected as biomarkers for CF patients, a plausible marker for cystic fibrosis, taking into account that the excess of PA available for ERAD translocation process must enter into turnover process.

In addition to PA and DG biomarkers, the present study has also identified an inflammatory mediator as CF biomarker: 19-(R)-hydroxy-PGF1α (407 *m/z*). The lack of functional CFTR in cystic fibrosis results in intrinsic inflammation, the major cause of morbidity and mortality in patients ^24a^. CFTR mutation induces the expression of high levels of nuclear factor kappa-light-chain-enhancer of activated B cells (NF-kappaB), and phospholipase A2 (PLA2) ^24b, 25^. In turn, these molecules induce the overproduction of inflammatory mediators, as well as cytokines and prostaglandins ^25-26a^, which is in accordance with the presented results. Furthermore, the redox imbalance caused by F508del-CFTR altered ion transportation causes abnormal generation of reactive oxygen species, leading to exacerbation of oxidative stress and development of inflammatory and degenerative lesions in target issues ^27^. Thereby, the increased inflammatory response and oxidative stress corroborate the election of hydroxylated prostaglandin F1-alpha as a CF biomarker.

Considering that mutant and misfolded proteins, as well as F508del-CFTR, are submitted to intense degradation through proteasome pathway, it is important to evaluate the products of amino acids catabolism, which will stand out from healthy cells metabolites. The conventional catabolism pathway of F508del-CFTR misfolded proteins occurs through glutaryl-CoA dehydrogenase and, alternatively, by the production of glutaric acid and its derivates, which justifies the election of 2,3-dimethyl-3-hydroxy-glutaric acid (211 *m/z*) as a CF biomarker. Considering that this biomarker is typically overproduced on ketogenic patients ^28a^, it is important to ponder that around 18.5% of the CFTR protein is composed by Lysine and Leucine, two purely ketogenic amino acids ^29^. Thus, the identification of 2,3-dimethyl-3-hydroxy-glutaric acid on CF skin is plausible, since most of F508del-CFTR misfolded proteins are under intense degradation process in proteasomes.

Given that F508delCFTR alters ionic transportation, many intracellular processes may be affected, including hypothalamic-pituitary-thyroid axis. Several studies have reported subclinical hypothyroidism on cystic fibrosis patients, and the absence of CFTR has been tested to elucidate its importance on synthesis of thyroid hormones ^30^. Within this context, Li and coauthors suggested that the increased Na^+^ absorption due to CFTR-/- may contribute to hypothyroidism on CF patients ^31^. Since CF patients may be affected by low levels of thyroid hormones, every cell responsible for its synthesis might be altered. In addition to the thyroid gland, skin cells are also responsible for the synthesis and metabolism of several hormones, including thyroid-stimulating hormone (TSH) ^32^. Cutaneous TSH expression is increased by thyrotropin-releasing hormone (TRH) and reduced by thyroid hormones ^32-33^. Since CF population is reported to present subclinical hypothyroidism and iodine deficiency ^34^, it is expected that high levels of TRH (397 *m/z*) be found in skin samples of CF patients. Interestingly, TRH was elected as a CF marker by the proposed methods in our study; this brings to our attention that a more frequent follow-up regarding the levels of thyroid hormones is necessary, since these tests are not routinely performed.

Another physiological important alteration in cystic fibrosis patients is CF liver disease (CFLD), a serious complication responsible for 2-4% of total mortality. The diagnosis of CFLD lack specific and sensitive markers; however, the prevalence rate ranges between 27 and 35% in patients older than 18 years ^35b^. Steatosis, for instance, can affect between 23% and 67% of CF patients of any age, presenting bile alkalization and viscosity increases. These events, in turn, result in higher levels of free radicals and accumulation of hydrophilic bile acids, which may provide damage to hepatocytes and bile ducts ^35a^. When in high concentration in the bloodstream, bile acids may be deposited on the skin, which corroborates the election of primary bile acids (425 *m/z* and 435 *m/z*) as CF markers. Taking into account that 67% of patients selected for this study were suspected of CFLD, the identification of bile acids among CF skin surface molecules is coherent with the evaluated patients. Although 44% of CF individuals evaluated in this study were under treatment with ursodeoxycholic acid, a drug used to reduce bile viscosity, its effectiveness has not been confirmed yet, and thus bile acids cannot be ruled out as potential biomarkers.

As shown above, the chosen markers are not only involved with the ionic imbalance caused by cystic fibrosis in the sweat gland, but also with the pathophysiology and progression of the disease; therefore, they may be considered relevant markers when monitored together for the F508del mutation. Although this contribution has evaluated only an F508del scenario, it is possible to infer that this methodology may also identify biomarkers in CF patients with any of the mutations, since the platform is sufficiently sensitive to detect small levels of altered components associated with physiological changes caused by other mutations. Nonetheless, further investigation is required, with a heterogeneous CF group, for conclusive results.

## 5. CONCLUSION

Taking into account the results shown above, the method applied for sample collection showed simplicity and less probability of methodological errors, differently from the standard method currently used. In addition, the proposed methodology using mass spectrometry analysis combined with statistical markers election allowed identifying the above-prospected biomarkers. When observed in a cystic fibrosis pathological condition, these biomarkers gain context and evidence points of metabolic disturbances that might be carefully evaluated as potential tools for new diagnostic investments. Interestingly, this contribution has evidenced the identification of primary bile acids as biomarkers for CF, which demonstrates that the concept of this approach may be useful not only for prospecting molecules that render new guides for diagnosis of CF, but may also be used for monitoring the progression of the disease. Therefore, the proposed method was capable of identifying potential cystic fibrosis diagnostic markers, in addition to evaluating disease evolution. These are valuable information for cystic fibrosis management and improvement in patients’ quality of life.

## ACKNOWLEDGEMENTS

We thank the Cystic Fibrosis Clinic Team – Clinical Hospital – University of Campinas for providing samples and support from the subjects involved in this research.

